# Quantifying Numerical and Spatial Reliability of Amygdala and Hippocampal Subdivisions in FreeSurfer

**DOI:** 10.1101/2020.06.12.149203

**Authors:** Isabella Kahhale, Nicholas J Buser, Christopher R. Madan, Jamie L. Hanson

## Abstract

On-going, large-scale neuroimaging initiatives can aid in uncovering neurobiological causes and correlates of poor mental health, disease pathology, and many other important conditions. As projects grow in scale with hundreds, even thousands, of individual participants and scans collected, quantification of brain structures by automated algorithms is becoming the only *truly* tractable approach. Here, we assessed the spatial and numerical reliability for newly deployed automated segmentation of hippocampal subfields and amygdala nuclei in FreeSurfer 7. In a sample of participants with repeated structural imaging scans (*N=923*), we found numerical reliability (as assessed by intraclass correlations, ICCs) was reasonable: ∼95% of hippocampal subfields have “excellent” numerical reliability (ICCs≥0.90), however, only 67% of amygdala subnuclei met this same threshold. Spatial reliability was similarly reasonable, with 58% of hippocampal subfields and 44% of amygdala subnuclei having Dice coefficients≥0.70. Notably, multiple regions had poor numerical and/or spatial reliability. We also examined correlations between spatial reliability and person-level factors (e.g., participant age; T1 image quality). Both sex and image scan quality were related to variations in spatial reliability metrics. Examined collectively, our work suggests caution should be exercised for a few hippocampal subfields and amygdala nuclei with more variable reliability.

## 1. Introduction

A large body of research has persistently emphasized that the hippocampus and amygdala play key roles in emotion and stress-responding. Critically, both of these subcortical structures show volumetric alterations in different neurodegenerative diseases and various forms of psychopathologies, including Alzheimer’s, Major Depression, Anxiety Disorders, and Autism (Campbell et al., 2004; Hamilton et al., 2008; Stanfield et al., 2008; Yang et al., 2012). Continued study of these regions could be critical in understanding cognitive processes—such as memory, decision making, emotion—and may lead to novel intervention strategies for different disorders.

Early studies focused on the hippocampus and amygdala typically examined volumes of these regions using expert manual tracing (Caldwell et al., 2015; Gunten et al., 2000; Hanson et al., 2015; MacQueen et al., 2003; Malykhin et al., 2008; von Gunten & Ron, 2004; Yucel et al., 2007). These approaches were at the time necessary to obtain reliable and valid measures of the size of these key brain areas, but hand-tracing is often exceedingly time intensive. As work in this space has continued, large-scale structural MRI-datasets (Ns from 100s to 1000s) are now more commonly available and work has shifted from manual tracing of regional volumes. Researchers are now able to leverage ever-improving computational algorithms to automatically segment structural images into their component anatomical structures (Madan & Kensinger, 2017). These approaches represent a scalable and less demanding method to test relations between volumetric measures of these two structures and psychological variables of interest.

A commonly used software suite, FreeSurfer (Fischl, 2012) provides a host of functions for structural MRI processing and analysis, including segmenting subcortical structures. Past work has examined both the validity and reliability of hippocampus and amygdala segmentation in FreeSurfer (Jovicich et al., 2009; Madan & Kensinger, 2017; Morey et al., 2010). One can think of validity as how well an output aligns with “ground-truth” (e.g., comparing FreeSurfer automated amygdala segments to expertly hand-traced volumes), while reliability reflects consistency of outputs (e.g., comparing FreeSurfer automated amygdala from repeated scans of the same person, without consideration of any “ground-truth”). Previous work has found strong reliability for FreeSurfer in terms of hippocampus and amygdala segmentations. Published reports examining test-retest reliability of subcortical volume measures have noted intraclass correlations from FreeSurfer ranging from 0.977-0.987 for the hippocampus and 0.806-0.889 for the amygdala (Jovicich et al., 2013; Liem et al., 2015; Wonderlick et al., 2009). Results considering validity have been more mixed. Work has investigated validity by comparing the spatial and numeric overlap between the volumes produced by FreeSurfer against those produced by expert hand tracing, finding reasonable Dice coefficients for the hippocampus, but lower performance on the amygdala (Hippocampus Dice coefficient=0.82; Amygdala Dice coefficient=0.72) (Hanson et al., 2012; Morey et al., 2009).

In considering both the hippocampus and amygdala, each of these brain areas are often discussed as unitary structures; however, a large body of basic molecular and cognitive neuroscience research underscores that the hippocampus and amygdala each consist of multiple distinct subregions with different information-processing roles. For example, the hippocampus can be subdivided into the following regions: Dentate Gyrus, critical for pattern separation (Neunuebel & Knierim, 2014); Cornu Ammonis (CA) 3, central to pattern completion (Guzman et al., 2016); CA 1, important for input integration from CA3 and entorhinal cortex (Bittner et al., 2015); and Subiculum, relevant for memory retrieval (Roy et al., 2017). Most of the past structural neuroimaging work has combined all these regions, using measures of whole hippocampal volume. This may mean a loss of specificity regarding associations with basic cognitive processes as well as neurobiological alterations seen in different disorders. By examining subcortical structure at a more fine-grain scale, results can be more precisely fit to their root cause and better interpreted considering their theoretical implications.

Responding to this issue, the developers of FreeSurfer have expanded their segmentation methods to include a more granular segmentation of hippocampal subregions (Iglesias et al., 2016). To do this, they combined ultra-high-resolution T1-weighted scans of post-mortem samples with subfields of the hippocampus segmented by hand, to develop an automated algorithm. With this algorithm, there appears to be good numerical reliability and slightly lower spatial reliability for these segments, mirroring the reliability work focusing on the whole hippocampus. Numerical reliability and ICCs are focused on the consistent overall volume size (as indexed by the number of voxels in a region), whereas spatial reliability and the calculation of Dice coefficients assess that the set of voxels classified are the same across both cases. These forms of reliability are typically correlated, but segments could have high numerical reliability but low spatial reliability. In such a case, the same number of voxels are being labelled as a brain region, but the voxels are in fact spatially divergent (and may not be the same brain area). Past work has observed high numerical and moderately high spatial reliability for the hippocampal subfields, reporting ICCs ranging from 0.70 to 0.97 and Dice coefficients ranging from approximately 0.60-0.90 (Brown et al., 2020; Whelan et al., 2016).

The amygdala, similarly, has its own subdivisions and the reliability of these subdivisions are still unclear. The FreeSurfer team also expanded their segmentation pipeline to cover a set of subdivisions for the amygdala. The algorithm they employ is trained on manually segmented amygdala nuclei from high-definition 7 Tesla ex-vivo MR images and divides this structure into 9 labelled sub-regions. They applied this segmentation to datasets looking at populations with autism (Di Martino et al., 2014) and those at risk for Alzheimer’s disease (Jack et al., 2008) finding significant improvements in pathology detection when this more fined grained view of the amygdala was used in the model (Saygin et al., 2017). However, direct assessment of numerical and spatial reliability for amygdala subdivisions is limited. Quattrini and colleagues (2020) examined these segments in a modest cohort of individuals (total *N=133*) and found reasonable reliability for larger subdivisions (>200 mm^3^ for the amygdala; >300 mm^3^ for the hippocampus). This work, however, aggregated across 17 research sites and multiple MRI vendors, deployed a dated version of the software (Freesurfer 6.0), and often acquired repeated imaging scans across weeks and months. Given these limitations, the consistency of these segments is an open question, and it is still unclear whether this fine-grained separation is consistent in the areas that the algorithm is automatically dividing and outputting. Such gaps are critical to fill given that many groups are using these algorithms for applied purposes and reporting differences between clinical and non-clinical populations (Morey et al., 2020; Zheng et al., 2019).

Motivated by these facts, we seek to provide an in-depth examination of reliability, both numerically and spatially, for FreeSurfer derived hippocampal and amygdala subdivisions. We leverage a public-access dataset of repeated structural scans with a large sample size (*N=928*). In addition to this first-order goal, we also wanted to consider whether person-level (e.g., age, sex) and MR-acquisition (e.g., MRI quality) factors influence the reliability of these subdivisions. Of note, recent work suggests that MR quality can significantly drive signal variations in structural MRI analyses (Gilmore et al., 2021; Madan & Kensinger, 2017). Pursuing these aims can inform whether all subdivisions are truly “reliable” and should be explored in FreeSurfer-related analyses, or if caution should be taken in morphometric comparisons (especially for those working in applied areas, e.g., tests of amygdala subdivisions in depressed vs. non-depressed groups).

## 2. Methods

### Participants

Data from an open-access neuroimaging initiative, the Amsterdam Open MRI Collection (AOMIC) (Snoek et al., 2021) was used to investigate numerical and spatial reliability of FreeSurfer’s amygdala and hippocampal subregion segmentation algorithms. AOMIC includes structural and functional neuroimaging scans from participants, repeating scans in the same session to see the stability of MRI-based metrics. For this work, data from 928 participants (Average Age=22.08, Standard Deviation =1.88) was examined. The majority of participants (n = 913, 98% of the sample) had three T1-weighted MR images collected in the same scanning session, while a small subgroup of participants (n = 15, ∼2% of the sample) had two T1-weighted scans. Additionally, 5 subjects were removed for extreme outlier values suggesting a processing error. The final analytic was 923 participants. All repeated MRI scans were acquired with the same imaging parameters (noted below).

### MRI Scan Parameters

MR images were acquired with a Phillips 3T scanner (“*Intera*” version) at the University of Amsterdam. T1-weighted MR images were acquired using a sagittal 3D-MPRAGE sequence (TR/TE = 8.1ms/3.7ms, 1mm^3^ voxel, matrix size = 64×64). Additional details about the scanning parameters are described by Snoek and colleagues (2021). MRI Images were visually inspected to determine if a participant’s scans should be included in subsequent processing steps (e.g., FreeSurfer).

### Structural Neuroimaging Processing (FreeSurfer)

Standard-processing approaches from FreeSurfer (e.g., cortical reconstruction; volumetric segmentation) were performed in version 7.1 (Stable Release, May 11, 2020) This was implemented via Brainlife.io (<u>brainlife.app.0</u>), a free, publicly funded, cloud-computing platform that allows for developing reproducible neuroimaging processing pipelines and sharing data (Avesani et al., 2019; Pestilli, 2018). FreeSurfer is a widely documented and freely available morphometric processing tool suite (http://surfer.nmr.mgh.harvard.edu). The technical details of this software suite are described in prior publications (Dale et al., 1999; Fischl, 2012; Fischl et al., 2002, 2004; Fischl, Sereno, & Dale, 1999; Fischl, Sereno, Tootell, et al., 1999). Briefly, this processing includes motion correction and intensity normalization of T1-weighted images, removal of non-brain tissue using a hybrid watershed/surface deformation procedure (Ségonne et al., 2004), automated Talairach transformation, segmentation of the subcortical white matter and deep gray matter volumetric structures (including hippocampus, amygdala, caudate, putamen, ventricles), tessellation of the gray matter white matter boundary, and derivation of cortical thickness. Scans from two subjects failed to run to completion in this pipeline and both subjects were removed from further analysis.

FreeSurfer version 7.1 natively includes options to segment hippocampal subfields and amygdalar nuclei. The hippocampal segmentation method (Iglesias et al., 2015) is based on a hippocampal atlas initially produced from a dataset of 15 hand-traced high definition ex-vivo T1-weighted 7 T scans then applied to a set of 39 standard resolution in-vivo MPRAGE scans using parameterized mesh deformations and a probabilistic atlas classification approach. This atlas is used for algorithmic segmentation of MR images pre-processed through the FreeSurfer recon-all pipeline. These images were classified using a parameterized generative model and optimizing the likelihood that any given voxel belongs to the label of a particular hippocampal region in a Bayesian inference framework (for additional information, see Iglesias et al., 2015). The atlas for this method partitions the hippocampus into the following 12 subfields: (1) Parasubiculum, (2) Presubiculum [Head and Body], (3) Subiculum [Head and Body], (4) CA1 [Head and Body], (5) CA3 [Head and Body], (6) CA4 [Head and Body], (7) Granule Cell and Molecular Layer of the Dentate Gyrus [GC-ML-DG, Head and Body], (8) Molecular layer [Head and Body], (9) Fimbria, (10) Hippocampal Fissure, (11) Hippocampal Tail, and (12) Hippocampus-Amygdala-Transition-Area (HATA). This yields nineteen subdivisions from FreeSurfer (including these regions and head/body divisions).

For the amygdala, the automated segmentation method (Saygin et al., 2017) is based on an atlas produced from 10 hand-traced high definition ex-vivo T1w 7 T scans (5 participants traced bilaterally). As in the hippocampal atlas, this manually segmented ex-vivo data was then applied to the probabilistic classification of the nodes on a parameterized deformation mesh of the amygdala. Similar to the hippocampus, the segmentation of later input data is performed in the framework of Bayesian inference. The amygdala atlas partitions the structure into the following 6 subnuclei: (1) Lateral, (2) Basal, (3) Central, (4) Medial, (5) Cortical, (6) Accessory Basal, (6) Paralaminar. Two additional subdivisions, the Corticoamygdaloid Transition Area and Anterior Amygdaloid Area, are also output.

Of note, here we processed two scans from each participant using the “*cross-sectional*” pipeline. This is in contrast to FreeSurfer’s longitudinal stream that creates an unbiased within-subject template image to improve temporal consistency and reduce potential source of bias (e.g., misregistration) (Reuter et al., 2010, 2012). Cross-sectional pipelines were applied to the three scans for each participant. For both the hippocampal subfields and amygdala nuclei, volume (in mm^3^) for each subdivision was extracted and used in numerical reliability analysis. Spatial information (labelled voxels in axial, coronal, and spatial orientations) was output for each subdivision. Each participant’s T1-weighted scan was then transformed to a common space using FMRIB’s Linear Image Registration Tool (degrees of freedom = 6; registering the 2^nd^ and 3^rd^ scans to the participant’s 1^st^ scan). This transformation matrix was then saved and applied to each volume’s labelled output for hippocampal and amygdala subdivisions using a nearest neighbour interpolation; these transformed hippocampal and amygdala subdivisions were then used in spatial reliability analysis.

### Automated MRI Image Quality Assessment

The Computational Anatomy Toolbox 12 (CAT12) toolbox from the Structural Brain Mapping group, implemented in SPM12, was used to generate a quantitative metric indicating the quality of each collected MR image (Gaser & Kurth, 2017). The method employed considers four summary measures of image quality: (1) noise to contrast ratio, (2) coefficient of joint variation, (3) inhomogeneity to contrast ratio, and (4) root mean squared voxel resolution. To produce a single aggregate metric that serves as an indicator of overall quality, this toolbox normalizes each measure and combines them using a kappa statistic-based framework, for optimizing a generalized linear model through solving least squares (Dahnke et al., 2015). After extracting one quality metric for each scan, we generated three values that represent the difference between two scans (i.e., Scan 1 – Scan 2; Scan 1 – Scan 3; Scan 2 – Scan 3). After taking the absolute value of each of these difference scores, we then averaged them together and used this as a measure of aggregate image quality.

### Derivation of Reliability Measures

To assess the reliability of numerical volumes output for hippocampus and amygdala subdivisions, we computed intraclass correlation coefficients (ICC) between each labelled sub-region for the test and the retest MRI scans. Of note, an ICC is a descriptive statistic indicating the degree of agreement between two (or more) sets of measurements. The statistic is similar to a bivariate correlation coefficient insofar as it has a range from 0-1 and higher values represent a stronger relationship. An ICC, however, differs from the bivariate correlation in that it works on groups of measurements and gives an indication of the numerical cohesion across the given groups (McGraw & Wong, 1996). The ICC was calculated separately for each sub-region using the statistical programming language R, with the icc function from the package *’irr’* (Gamer & Lemon, 2012). A two-way model with absolute agreement was used in order to investigate the reliability of subdivision segmentation; this was calculated for each subdivision’s volume (in mm^3^). Although there are no definitive guidelines for precise interpretation of ICCs, results have frequently been binned into three (or four) quality groups where 0.0-0.5 is “poor”, 0.50-0.75 is “moderate”, 0.75-0.9 is “good” and 0.9-1.0 is “excellent” (Cicchetti, 1994; Koo & Li, 2016).

In addition to ICCs, Bland-Altman metrics were calculated for each hippocampal and amygdala subdivision using the function blandr.statistics from the package ’*blandr’* (Datta, 2017). In this approach, the mean differences (“*bias*”) between the FreeSurfer outputs (comparing the first and second scan, the first and third scan, and the second and third scan) were first calculated and presented as a portion of the mean volume. We took the absolute value of each of these three values and averaged them together to represent the average Bland-Altman metric across the three scans for a given brain region. Bland-Altman plots were also constructed for a small number of subdivisions to assess agreement between FreeSurfer outputs.

Although ICCs and Bland-Altman metrics serve as indicators of numerical reliability, these may still be incomplete, particularly when we think about the spatial information present in MRI volumes. Indeed, even with numerical similarity, there may be discrepancies in the specific spatial voxels labelled for a given subdivision. To assess whether the voxels assigned to each region were the same between the two timepoints, we calculated the Sørensen-Dice Coefficient using the @DiceMetric program in the AFNI fMRI software package (Cox, 1996). The Dice coefficient is calculated as (2TP)/(2TP + FP + FN) [TP = True Positive; FP = False Positive; FN = False Negative] and gives an equal weight to criteria of positive predictive value and sensitivity in assessing spatial reliability of subdivisions. Dice coefficients were averaged across the three scans for each brain region to obtain an overall metric of spatial reliability (e.g., one Dice value for the Left Lateral Nucleus, one Dice value for the Right Lateral Nucleus). As recommended by past reports (Zijdenbos et al., 1994; Zou et al., 2004), we considered Dice Coefficients ≥0.700 as exhibiting “good” spatial overlap.

### Statistical Analysis

Once overall reliability metrics were calculated, we examined person-level (e.g., age, sex) and MR-acquisition (e.g., MRI quality) factors in relation to these measures. Many different factors may impact amygdala and hippocampal segmentation. For example, past work suggests volumes (of the hippocampus and amygdala) vary with participant age and sex; this association is particularly strong for the hippocampus (Daugherty et al., 2016; Nobis et al., 2019) and suggestive data similarly for the amygdala (Malykhin et al., 2008; Marwha et al., 2017; Perlaki et al., 2014; Pressman et al., 2016). Finally, image quality has been shown to have a significant effect on brain volume measurements (Gilmore et al., 2021). Noisier images may lead to gray/white matter misclassification, and impact reliability between different scans. To consider these potential effects, we examined each region’s reliability in relation to age, sex, and difference in the CAT12 quality metric. We constructed regression models to examine connections between reliability metrics, and person-level and MR-acquisition factors. Of note, the average difference in quality between the three scans (described in *Automated MRI Image Quality Assessment*) was included in these analyses.

## 3. Results

### Hippocampus Reliability

Using ICC analysis, we found consistently reasonable levels of numerical reliability for hippocampal subfields. Multiple regions demonstrated “excellent” reliability (ICC ≥ 0.90), while all of the subfields were at least in the “good” range (ICC=0.75-0.90). See Table 1 for values from the 19 subfield segmentations in each hemisphere. Bland-Altman bias indicated some variability with differences between scans, as a portion of that structure’s volume, ranging from 0.078-1.198%. See Figure 1 for a density plot of the average difference in volume estimation across three scans for two hippocampal subfields.

**Figure 1:**
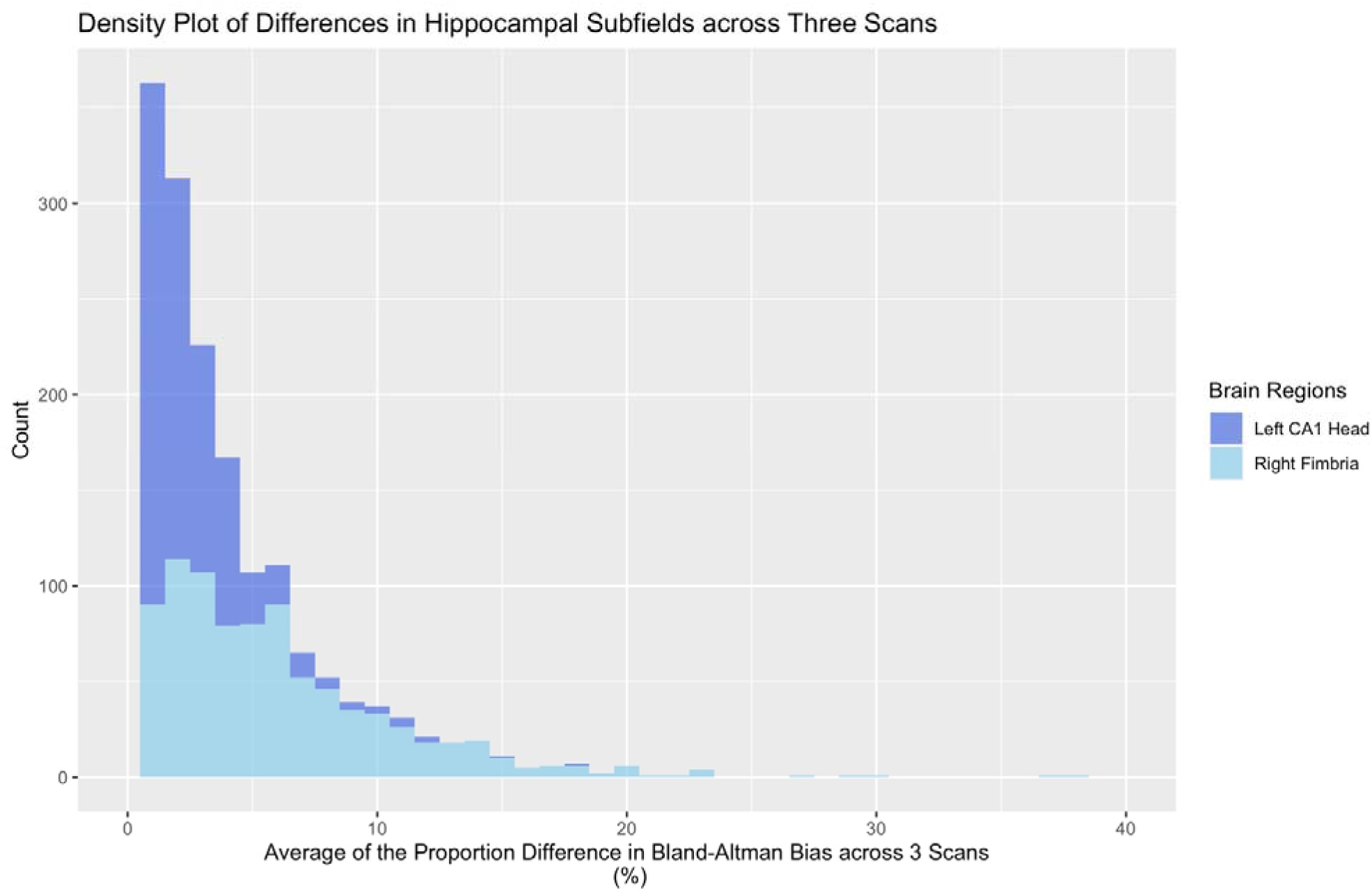
Bland–Altman plots of the average difference for volume estimation across subjects’ three MRI scans for the Left Cornu Ammonis (CA) 1 Head (dark blue) and Right Fimbria (light blue). The horizontal axis indicates the average difference in Bland-Altman “*bias*” (difference between subregional volume output for different scans, as a proportion of a region’s volume), while the vertical axis indicates the number of scans with a given value. Of note, the left CA1 Head has a low degree of mean bias (as a proportion of the region’s volumes; 0.106%), while the right Fimbria has a fair degree of mean bias (1.198%).

**Table 1:** Intraclass Correlation Coefficients (ICC), Dice Coefficients, Bland–Altman bias as a portion of a volume’s structure (Bias as POV), and Bland-Altman bias ranges for Hippocampal Subfields for left and right hemisphere regions (e.g., ICC LH = Intraclass Correlation Coefficients for left hemisphere; Dice RH = Dice Coefficient for right hemisphere). Color coding is in accordance with excellent [green], good [yellow], poor [red] scores for ICCs and Dice Coefficients (ICC: 0.90-1.00 [excellent], 0.75-0.89 [good], 0.00-0.74 [poor]; Dice Coefficients: 0.70-1.00 [excellent], 0.50-.69 [good], 0-0.49 [poor]). We have also highlighted regions with >1% bias as a portion of a volume’s structure in yellow. Subfield Abbreviations include: Cornu Ammonis (CA), Granule Cell and Molecular Layer of Dentate Gyrus (GC-ML-DG); Hippocampus-Amygdala-Transition-Area (HATA); Hippocampal Parcellation (HP).

| Region | ICC<br>LH | ICC<br>RH | Dice<br>LH | Dice<br>RH | Bias as<br>POV LH | Bias Range<br>LH | Bias as<br>POV RH | Bias Range<br>RH |
| --- | --- | --- | --- | --- | --- | --- | --- | --- |
| Parasubiculum | 0.929 | 0.946 | 0.713 | 0.710 | 0.356% | 0.000 – 45.834% | 0.410% | 0.008 – 32.920% |
| Presubiculum Head | 0.924 | 0.936 | 0.793 | 0.791 | 0.212% | 0.001 – 29.992% | 0.078% | 0.002 – 23.345% |
| Presubiculum Body | 0.960 | 0.963 | 0.800 | 0.791 | 1.057% | 0.000 – 41.021% | 0.626% | 0.001 – 33.210% |
| Subiculum Head | 0.961 | 0.959 | 0.777 | 0.773 | 0.130% | 0.001 – 27.278% | 0.085% | 0.002 – 25.029% |
| Subiculum Body | 0.961 | 0.964 | 0.823 | 0.826 | 0.100% | 0.004 – 31.485% | 0.289% | 0.002 – 16.760% |
| CA1 Head | 0.971 | 0.979 | 0.819 | 0.825 | 0.106% | 0.000 – 22.307% | 0.260% | 0.000 – 16.274% |
| CA1 Body | 0.948 | 0.970 | 0.752 | 0.780 | 0.127% | 0.001 – 50.541% | 0.465% | 0.001 – 29.594% |
| CA3 Head | 0.952 | 0.969 | 0.663 | 0.677 | 0.533% | 0.000 – 24.448% | 0.742% | 0.000 – 22.072% |
| CA3 Body | 0.933 | 0.950 | 0.598 | 0.627 | 0.413% | 0.000 – 55.834% | 0.973% | 0.001 – 27.936% |
| CA4 Head | 0.966 | 0.966 | 0.794 | 0.798 | 0.551% | 0.000 – 20.589% | 0.494% | 0.000 – 21.287% |
| CA4 Body | 0.931 | 0.938 | 0.768 | 0.780 | 0.541% | 0.002 – 29.764% | 0.625% | 0.002 – 20.289% |
| GC ML DG Head | 0.965 | 0.973 | 0.618 | 0.626 | 0.554% | 0.001 – 20.329% | 0.542% | 0.002 – 18.378% |
| GC ML DG Body | 0.940 | 0.942 | 0.593 | 0.600 | 0.500% | 0.000 – 21.930% | 0.503% | 0.001 – 27.053% |
| Molecular Layer HP Head | 0.971 | 0.976 | 0.690 | 0.692 | 0.206% | 0.001 – 18.600% | 0.240% | 0.001 – 14.279% |
| Molecular Layer HP Body | 0.954 | 0.961 | 0.632 | 0.647 | 0.495% | 0.001 – 25.208% | 0.558% | 0.000 – 18.751% |
| Fimbria | 0.936 | 0.942 | 0.681 | 0.679 | 0.639% | 0.008 – 78.157% | 1.198% | 0.004 – 57.572% |
| Hippocampal Fissure | 0.887 | 0.888 | 0.498 | 0.511 | 0.443% | 0.001 – 41.512% | 0.586% | 0.000 – 45.252% |
| Hippocampal Tail | 0.954 | 0.968 | 0.882 | 0.890 | 0.315% | 0.000 – 50.169 % | 0.393% | 0.001 – 18.664% |
| HATA | 0.936 | 0.945 | 0.760 | 0.770 | 1.049% | 0.000 – 34.712% | 0.216% | 0.007 – 26.125% |
| Whole Hippocampal Body | 0.959 | 0.966 |  |  | 0.365% | 0.000 – 27.935% | 0.435% | 0.004 – 15.245% |
| Whole Hippocampal Head | 0.975 | 0.980 |  |  | 0.227% | 0.001 – 19.277% | 0.273% | 0.002 – 13.619% |
| Whole Hippocampus | 0.976 | 0.983 |  |  | 0.251% | 0.001 – 21.668% | 0.341% | 0.000 – 12.473% |

Using Dice coefficients as metrics of spatial reliability, results became a bit more variable with 11 areas showing “excellent” spatial reliability, 7 areas showing “good” spatial reliability, and one area (left Hippocampal fissure) showing poor spatial reliability (Dice Coefficient<0.5.). See Figure 2 for a plot of all Hippocampal Dice Coefficient values and Figure 3 for an example of regions with acceptable spatial reliability (parasubiculum) and poor spatial reliability (hippocampal fissure).

**Figure 2.**
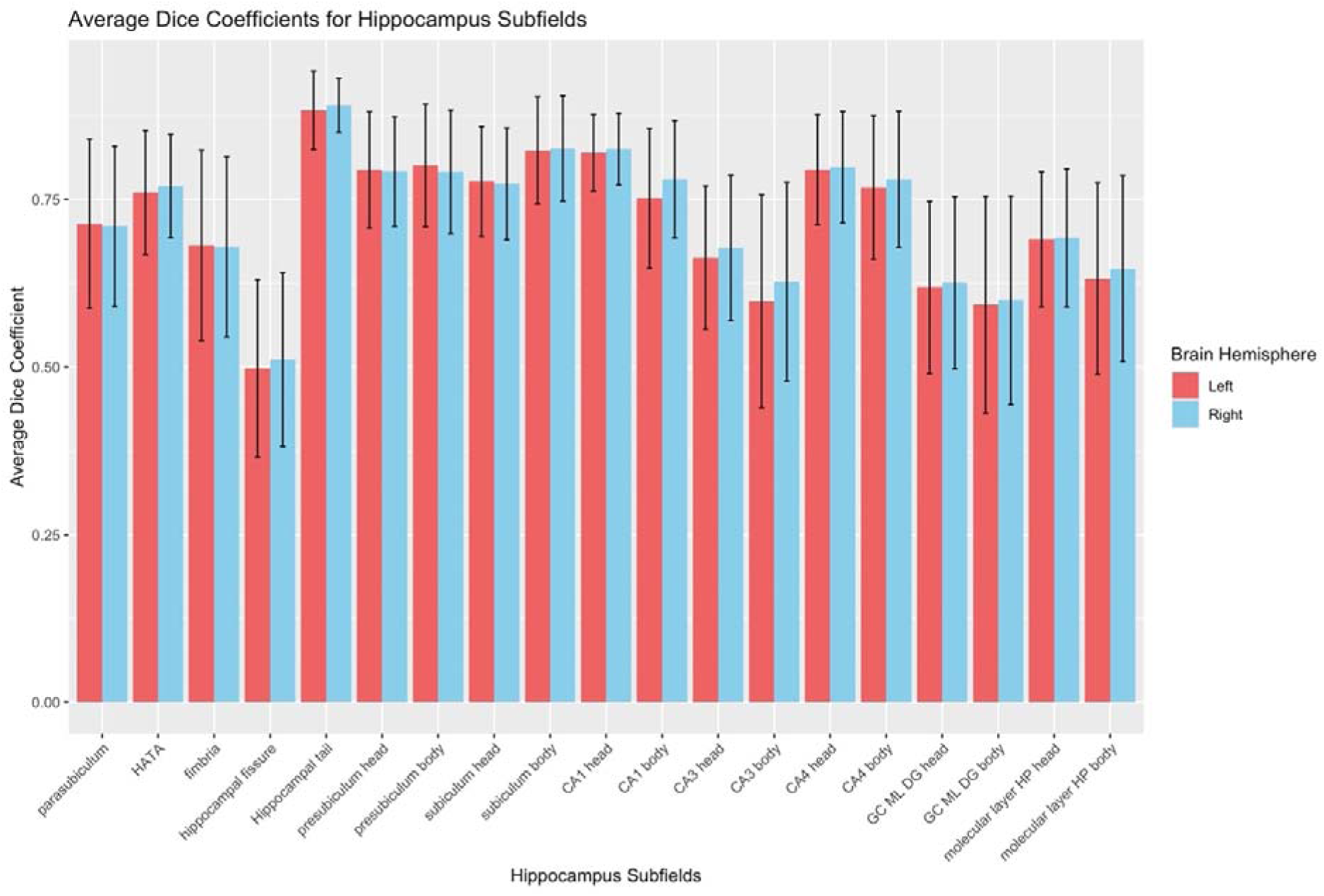
Hippocampal Dice Coefficient values for all hippocampal subfields. Error bars represent 1 standard deviation above and below the mean. Subfield Abbreviations include: Cornu Ammonis (CA), Granule Cell and Molecular Layer of Dentate Gyrus (GC-ML-DG); Hippocampus-Amygdala-Transition-Area (HATA); Hippocampal Parcellation (HP).

**Figure 3:**
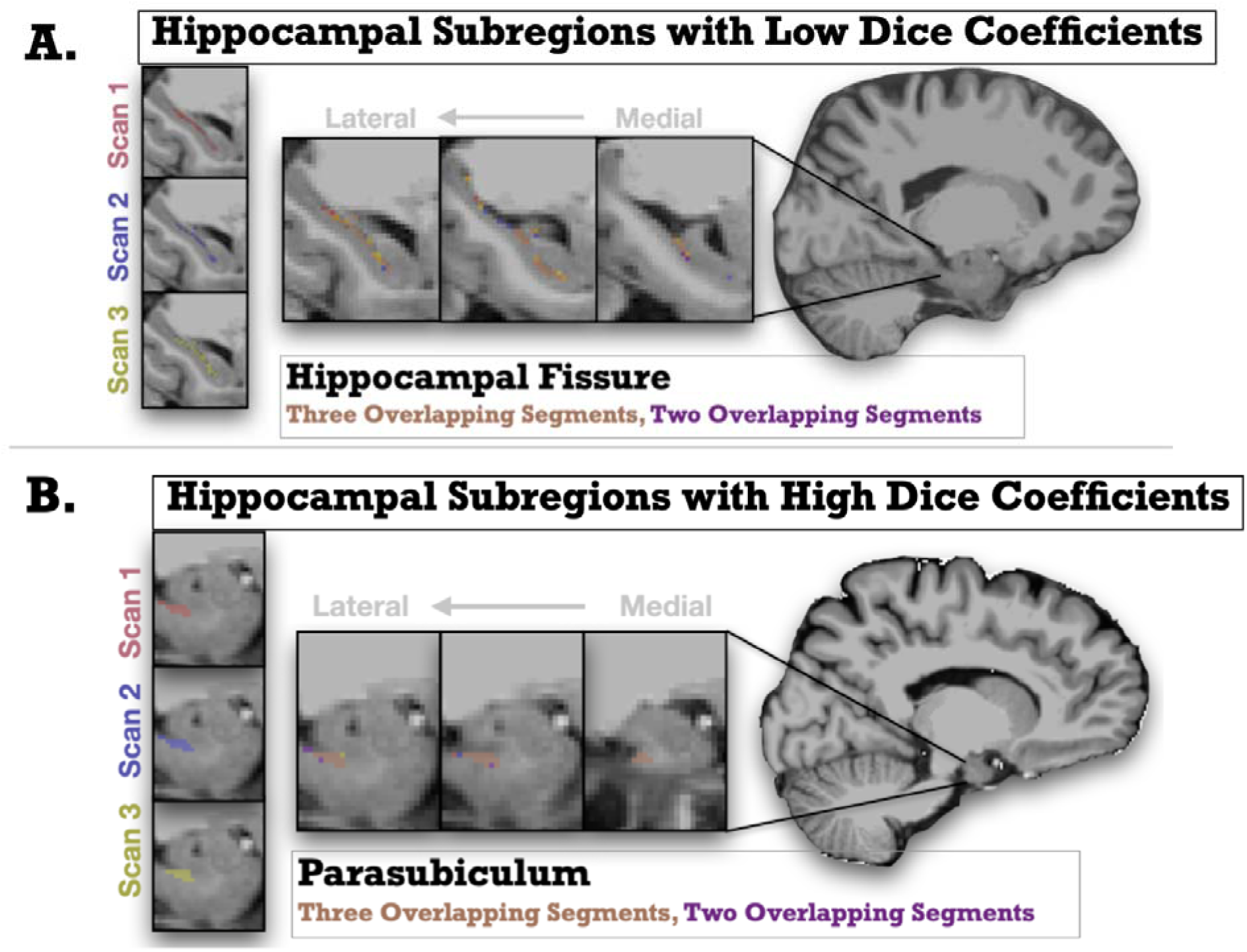
Graphic representations showing magnified depictions of Hippocampal subregions with low and high Dice coefficients (i.e., spatial reliability) from repeated scans (Scan 1 shown in Red; Scan 2 shown in Purple, Scan 3 shown in Yellow). The anatomical (T1w) image underlaid is the unbiased subject template from an example participant. The top panel A represents the hippocampal fissure, an area with low spatial reliability across scans, and the bottom panel B represents the parsubiculum, and area with high spatial reliability. Slices move right to left from medial to lateral.

### Amygdala Reliability

Within the amygdala, the numerical reliability was “excellent” for about 67% of the regions (ICC > .90), while the remainder of the regions were in the “good” range (ICC = 0.75-0.90) (*see Table 2*). Bland-Altman bias values were somewhat variable with a range of 0.058-1.563%. See Figure 4 for a density plot of the average difference in volume estimation across three scans for two amygdala subnuclei.

**Figure 4:**
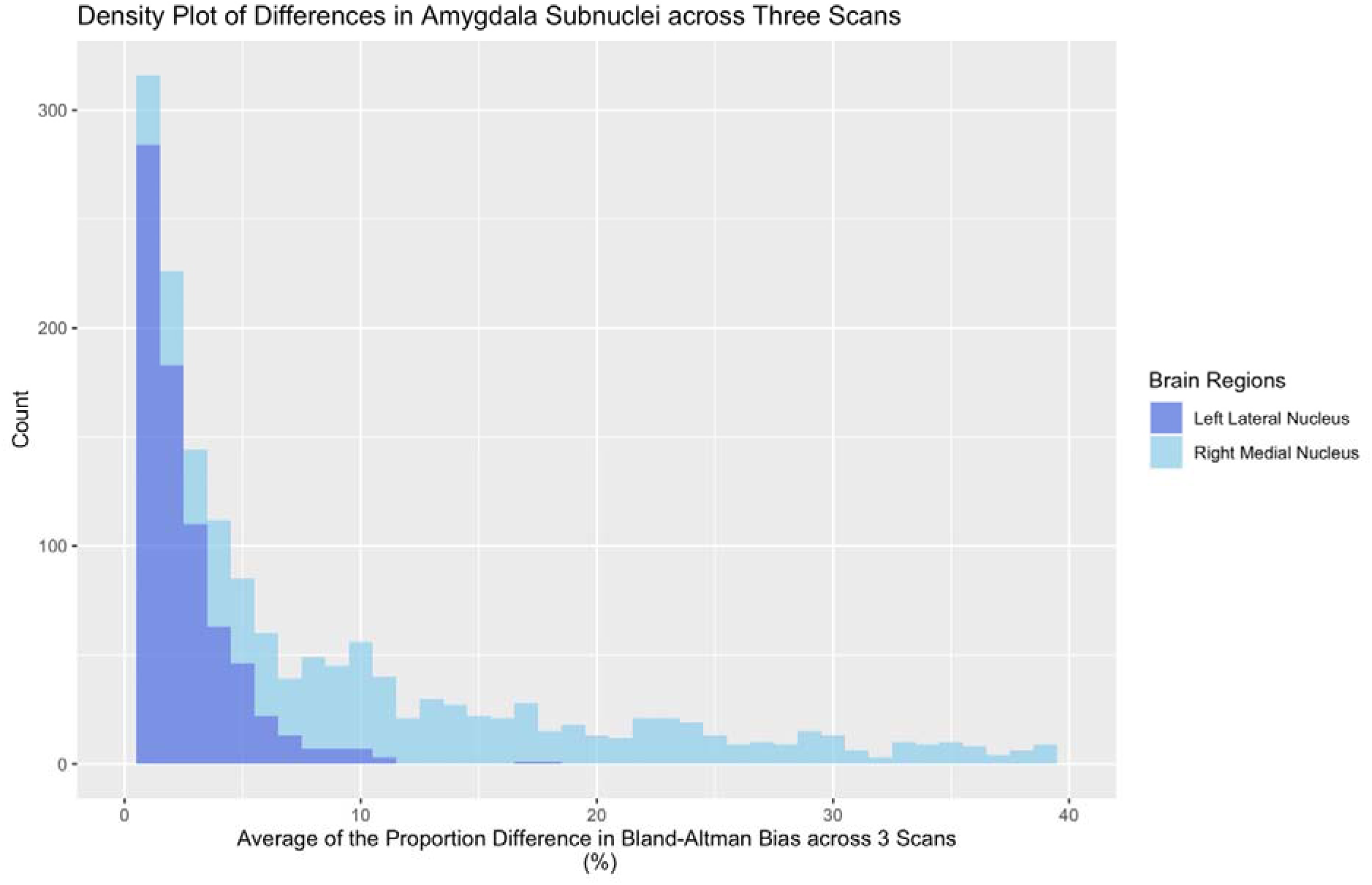
Bland–Altman plots of the average volume difference estimation across Scan 1, Scan 2, and Scan 3 for the left Lateral Nucleus (dark blue) and right Medial Nucleus (light blue). The horizontal axis indicates the average difference in Bland-Altman “*bias*” (difference between subregional volume output for different scans, as a proportion of a region’s volume), while the vertical axis indicates the number of scans with a given value. Of note, the left Lateral Nucleus has a low degree of bias (as a proportion of the region’s volumes; 0.108%), while the right Medial Nucleus has a fair degree of bias (1.047%).

**Table 2:**
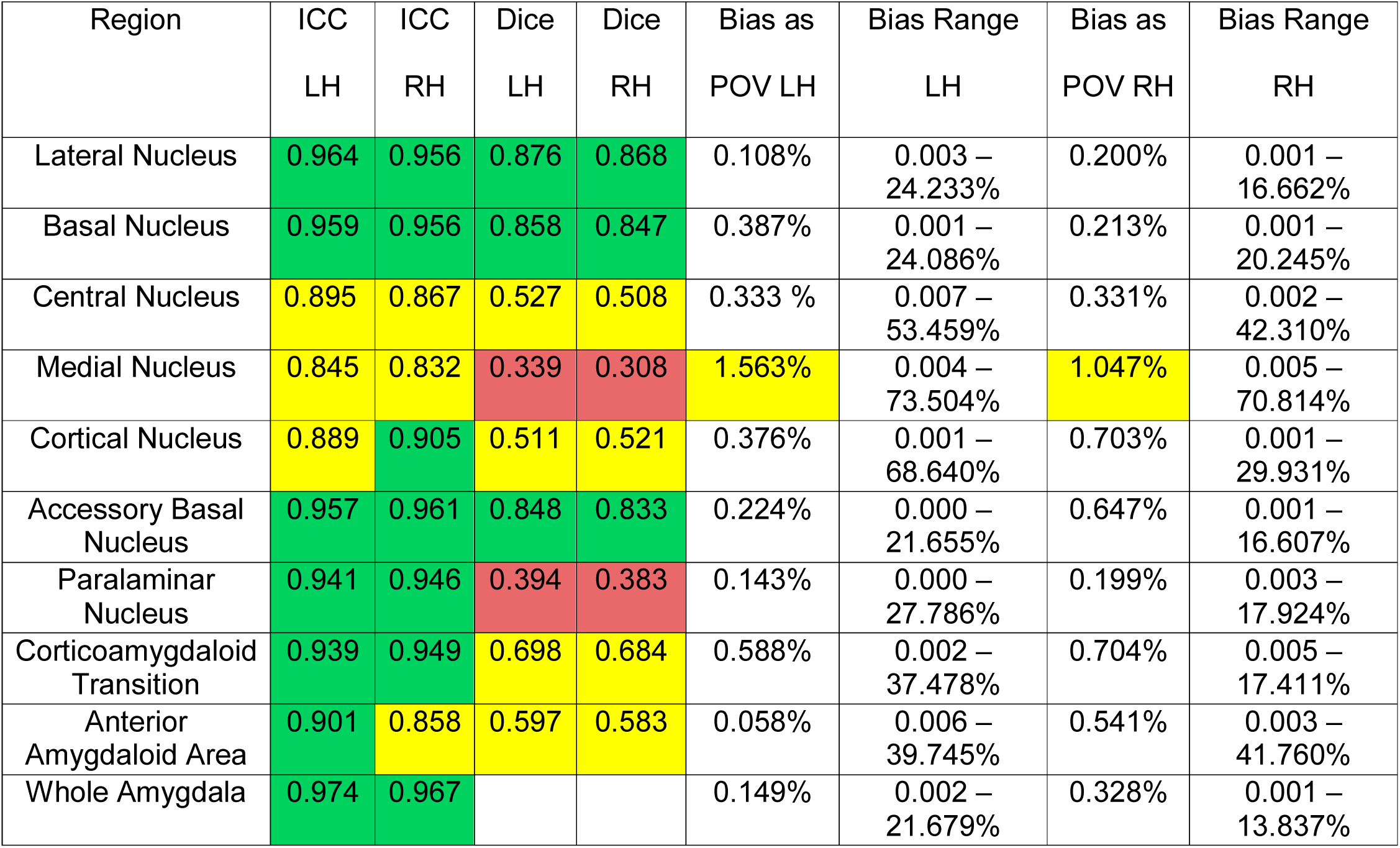
Intraclass Correlation Coefficients (ICCs), Dice Coefficients, Bland–Altman bias as a portion of a volume’s structure (Bias as POV), and Bland-Altman bias ranges for Amygdala Subnuclei for left and right hemisphere regions (e.g., ICC LH = Intraclass Correlation Coefficients for left hemisphere; Dice RH = Dice Coefficient for right hemisphere). Color coding is in accordance with excellent [green], good [yellow], poor [red] scores for ICCs and Dice Coefficients (ICC: 0.90-1.00 [excellent], 0.75-0.89 [good], 0.00-0.74 [poor]; Dice Coefficients: 0.70-1.00 [excellent], 0.50-.69 [good], 0-0.49 [poor]). We have also highlighted regions with >1% bias as a portion of a volume’s structure in yellow.

Regarding spatial reliability, several areas demonstrated good reliability (≥.7) including the lateral, basal, and accessory basal subnuclei (See *Table 2*). There were, however, areas with poor spatial reliability (Dice Coefficients = 0.30-0.4, See *Figure 5* for a plot of all Amygdala Dice Coefficient values). These problems were primarily seen in the Medial and Paralaminar Nuclei. Figure 6 displays a depiction of the Lateral nucleus, an area with acceptable spatial reliability, and the Paralaminar nucleus, an area with poor spatial reliability.

**Figure 5.**
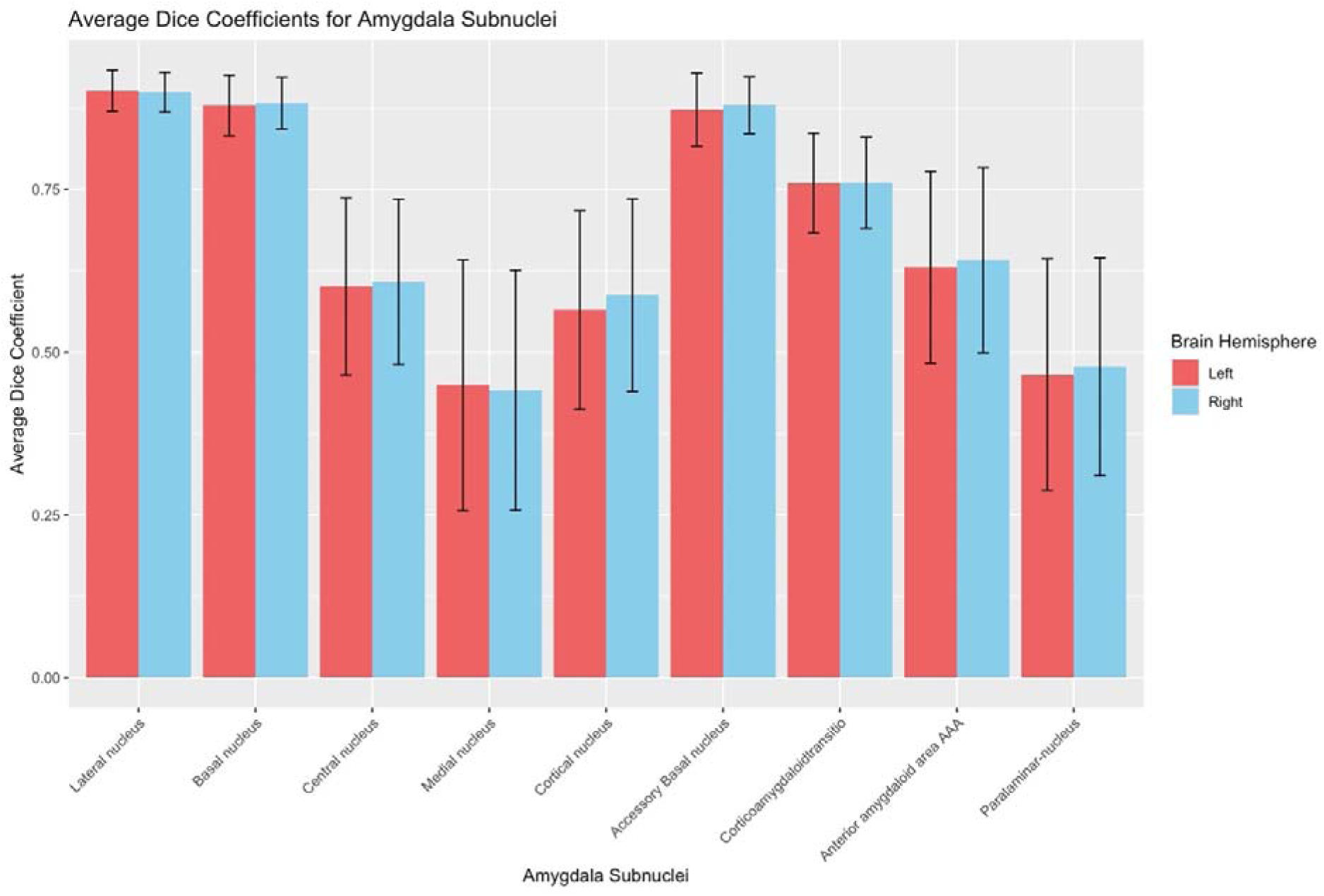
Amygdala Dice Coefficient values for all Amygdala subnuclei. Error bars represent 1 standard deviation above and below the mean.

**Figure 6:**
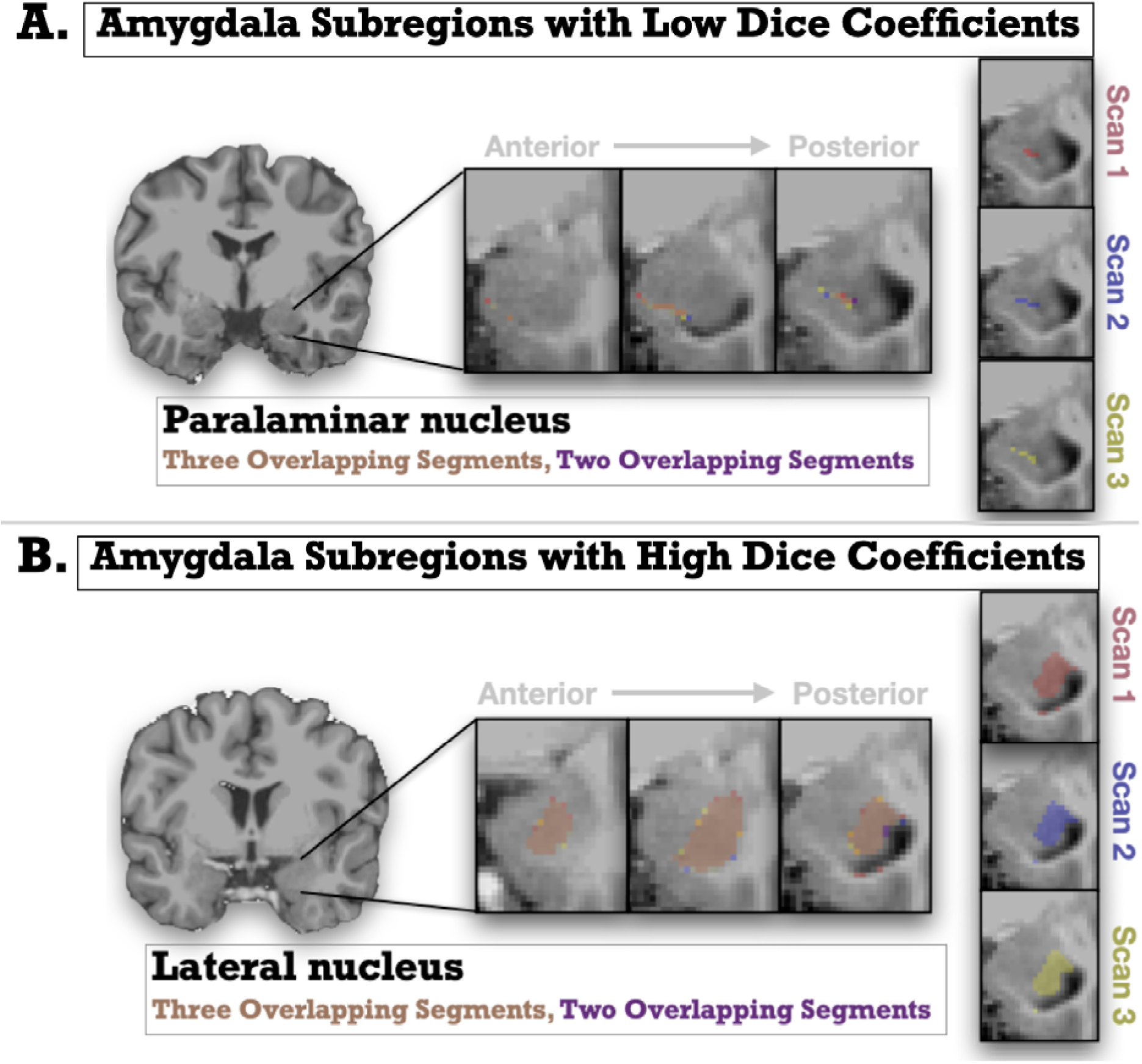
Graphic representations showing magnified depictions of Amygdala subregions with low and high Dice coefficients (i.e., spatial reliability) from repeated scans (Scan 1 shown in Red; Scan 2 shown in Purple, Scan 3 shown in Yellow). The anatomical (T1w) image underlaid is the unbiased subject template from an example participant. The top panel A represents the paralaminar nucleus, an area with low spatial reliability across scans, and the bottom panel B represents the lateral nucleus, and area with high spatial reliability. Slices move right to left from medial to lateral. Multiple slices are depicted left to right, moving anterior to posterior.

### Reliability Differences in Relation to Person-level and MR-acquisition Factors

We next examined associations between spatial reliability and subject-level variables. Correlations between the Hippocampal-subfield Dice coefficients and our subject-level variables are shown in Table 3. Differences in image quality and participant sex were significantly and negatively related to volumes in a majority of the hippocampal subfields at the p < 0.01 level (shown in Table 3). Age was not significantly correlated with any hippocampal subfield volumes.

**Table 3:** Correlation coefficient for bivariate correlations between Hippocampal Subfield Dice Coefficients and subject-level covariates: MRI Quality (Difference Score; MRIQ), Sex, and Age. These were completed for left hemisphere (LH) and right hemisphere (RH). Correlations with p<.01 are highlighted in yellow. Subfield Abbreviations include: Cornu Ammonis (CA), Granule Cell and Molecular Layer of Dentate Gyrus (GC-ML-DG); Hippocampus-Amygdala-Transition-Area (HATA); Hippocampal Parcellation (HP).

| Region | MRIQ r Dice |  | Sex r Dice |  | Age r Dice |  |
| --- | --- | --- | --- | --- | --- | --- |
|  | LH | RH | LH | RH | LH | RH |
| Parasubiculum | -0.13 | -0.08 | -0.03 | -0.09 | 0.00 | -0.05 |
| Presubiculum Head | -0.13 | -0.16 | -0.06 | -0.10 | -0.04 | -0.04 |
| Presubiculum Body | -0.15 | -0.16 | -0.08 | -0.08 | -0.02 | 0.01 |
| Subiculum Head | -0.13 | -0.17 | -0.06 | -0.09 | -0.03 | -0.05 |
| Subiculum Body | -0.09 | -0.11 | -0.06 | -0.09 | 0.02 | -0.03 |
| CA1 Head | -0.13 | -0.14 | -0.07 | -0.12 | -0.06 | -0.05 |
| CA1 Body | -0.04 | -0.07 | -0.08 | -0.13 | 0.01 | -0.06 |
| CA3 Head | -0.09 | -0.07 | -0.09 | -0.12 | 0.01 | -0.06 |
| CA3 Body | -0.05 | -0.08 | -0.09 | -0.10 | 0.01 | -0.01 |
| CA4 Head | -0.09 | -0.13 | -0.08 | -0.09 | -0.04 | -0.07 |
| CA4 Body | -0.04 | -0.11 | -0.09 | -0.08 | 0.01 | -0.04 |
| GC ML DG Head | -0.10 | -0.11 | -0.07 | -0.10 | -0.03 | -0.08 |
| GC ML DG Body | -0.06 | -0.10 | -0.09 | -0.10 | 0.01 | -0.05 |
| Molecular Layer HP Head | -0.13 | -0.14 | -0.06 | -0.11 | -0.05 | -0.07 |
| Molecular Layer HP Body | -0.07 | -0.10 | -0.08 | -0.10 | 0.02 | -0.05 |
| Fimbria | -0.12 | -0.17 | -0.01 | -0.05 | 0.02 | -0.02 |
| Hippocampal Fissure | -0.10 | -0.17 | -0.08 | -0.13 | -0.04 | -0.05 |
| Hippocampal Tail | -0.08 | -0.14 | -0.12 | -0.11 | -0.03 | -0.01 |
| HATA | -0.08 | -0.10 | -0.09 | -0.12 | 0.00 | -0.04 |

Correlations between spatial reliability and subject-level variables for the amygdala nuclei are reported in Table 4. Image quality was significantly and negatively related to a small number of regions including the lateral nucleus, the right basal nucleus, the corticoamygdaloid transition, and the right anterior amygdaloid area (at *p<0.01*).

**Table 4:** Correlation coefficient for bivariate correlations between Hippocampal Subfield Dice Coefficients and subject-level covariates: MRI Quality (Difference Score; MRIQ), Sex, and Age. These were completed for left hemisphere (LH) and right hemisphere (RH). Correlations with p<.01 are highlighted in yellow.

| Region | MRIQ r Dice |  | Sex r Dice |  | Age r Dice |  |
| --- | --- | --- | --- | --- | --- | --- |
|  | LH | RH | LH | RH | LH | RH |
| Lateral Nucleus | -0.12 | -0.18 | -0.10 | -0.11 | 0.01 | -0.03 |
| Basal Nucleus | -0.07 | -0.10 | -0.05 | -0.06 | -0.03 | -0.07 |
| Central Nucleus | -0.05 | 0.01 | 0.04 | 0.01 | -0.03 | -0.10 |
| Medial Nucleus | -0.07 | -0.01 | -0.09 | -0.06 | 0.00 | -0.01 |
| Cortical Nucleus | 0.00 | -0.04 | -0.06 | -0.04 | 0.04 | -0.02 |
| Accessory Basal Nucleus | -0.05 | -0.07 | -0.06 | -0.03 | -0.03 | -0.08 |
| Paralamina Nucleus | -0.03 | -0.07 | -0.01 | -0.04 | -0.02 | -0.02 |
| Corticoamygdaloid Transition | -0.12 | -0.10 | -0.08 | -0.07 | -0.02 | -0.06 |
| Anterior Amygdaloid Area | -0.07 | -0.13 | -0.09 | -0.06 | -0.02 | -0.05 |

The spatial reliability of a minority of regions was also significantly and associated with sex and age.

## 4. Discussion

In this paper, we assessed the numerical and spatial reliability of FreeSurfer’s hippocampal and amygdala subdivision segmentation algorithms. The ICC’s, serving as our indicator of numerical reliability, were reasonable (hippocampal subfields: 0.887-0.979; amygdala nuclei: 0.832-0.964), indicating that FreeSurfer is generally numerically reliable in providing overall volume for each subregion. Using Bland–Altman metrics of bias as an additional proxy of numerical reliability suggests a few regions exhibited a good bit of variability in segmentation from one scan to the next; specifically, 5 regions across the hippocampus and amygdala showed ≥1% bias in volume from one scan to the next. This is concerning given that individuals with dementia (i.e., Alzheimer’s disease) or recurrent mental health issues (i.e., depression) often only differ 1-5% from control groups in subcortical volumes (e.g., Jack et al., 2005; Logue et al., 2018; Schmaal et al., 2016). The Dice coefficients, serving as our indicator of spatial reliability, were reasonable, though lower than the ICCs. Of potential concern, a few subdivisions in both the hippocampus and amygdala had fairly low spatial reliability, suggesting unreliable segmentation. Examined collectively, applied researchers should take care when applying these types of automated segmentation techniques, especially if not thoroughly trained in amygdala and hippocampal anatomy.

While our results suggest that many of the volumetric outputs of amygdala and hippocampal subdivisions are mostly numerically reliable, the drop in spatial reliability may mean researchers should exercise caution in the analysis and interpretation of areas with poor spatial reliability. For example, the hippocampal fissure, paralaminar nucleus (amygdala), medial nucleus (amygdala) showed poor spatial reliability (≤.5) through their Dice coefficients. Because the spatial reliability of these areas is relatively poor, studies that interpret changes in volume within or across subjects might be using segmentations which contain improperly (or inconsistently) classified voxels within those regions. For example, several studies have already reported significant findings from the paralaminar nucleus of the amygdala (Morey et al., 2020; Zheng et al., 2019); given the questionable reproducibility of its anatomical bounding, these findings may require further verification.

Connected to spatial reliability, there are a few potential drivers of the substandard performance in this domain. First, these areas are fairly small and may be difficult to isolate. In such cases, even a few mislabelled voxels can greatly influence spatial overlap. Many of the areas with the lowest spatial reliability are also the smallest subdivisions. For example, the Paralaminar and the medial nuclei of the amygdala range between 20-60 mm^3^ in our sample and have some of the lowest spatial reliability values. However, and of note, this is not the only factor hampering performance, as other structures (of similar sizes) have reasonable spatial reliability values (e.g., HATA ≥ 0.765 Dice Coefficients; Parasubiculum ≥ 0.712 Dice Coefficients), while comparatively larger structures (e.g., hippocampal-fissure; CA3-body) demonstrate lower spatial reliability. Second, irregular MR contrast is often common to these areas, especially for the amygdala. Given the close vicinity to bone and sinuses, there is typically susceptibility-related dropout, field inhomogeneities and physiological artifacts in the amygdala and the hippocampus (Merboldt et al., 2001; Robinson et al., 2004; Windischberger et al., 2002). This may introduce inconsistent gray/white matter contrast, complicating isolation of different subdivisions. Finally, several amygdala and hippocampal subdivisions are irregularly and complexly shaped. For example, both the anterior and posterior borders of the amygdala are difficult to consistently demarcate (Achten et al., 1998; Convit, 1999; Watson et al., 1992, as cited in Entis et al., 2012). Many past reports using manual tracing actually employ “heuristics” rather than clear anatomical boundaries (e.g., Nacewicz et al., 2006 used a “semicircle substitution”).

Given these challenges, it will be important to think about novel approaches to break down the amygdala and hippocampus into smaller subdivisions, while still maintaining high validity and reproducibility. In regard to the hippocampus, there has been a great deal of progress made by the Hippocampal Subfields Group (https://www.hippocampalsubfields.com/); this is a collaboration of >200 imaging and anatomy experts worldwide that has established guidelines for appropriate MRI acquisition for researchers interested in the hippocampus, as well as developing candidate protocols for the segmentation of hippocampal subregions (eg., Olsen et al., 2019; Wisse et al., 2017; Yushkevich et al., 2015). This and other related work have suggested important ways to validate automatic segmentation, including not only comparison to manual delineations, but also replicating known disease effects (e.g., Mueller et al., 2018). Similar joint efforts are not, to our knowledge, currently underway for amygdala subnuclei segmentation. Convening such a collaborative could be particularly powerful moving forward, especially as debate has been fairly continuous regarding subdivisions of the amygdala at the histological level (e.g., Swanson & Petrovich, 1998). In the interim, or in the absence of that type of group, it may be reasonable to only consider more macro-level amygdala segmentation (e.g., basolateral, centromedial, basomedial, and amygdaloid cortical complexes, as detailed by (Mai et al., 2015). Many groups have moved towards this idea, aggregating subdivisions using latent factor modelling and other techniques to group related regions (e.g., Oshri et al., 2019). There is, however, ongoing debate about specific best practices, as even established guidelines for MRI acquisition or landmark in in-vivo data may present additional unforeseen challenges (e.g., Special hippocampal acquisitions providing incomplete coverage of target structure; In-vivo MRI does not supply enough features to define many hippocampal subfield boundaries).

Of importance and as noted in our introduction, our findings only speak to the reliability of these measures, and not the validity of these segments. Future work should be pursued to establish the validity of these FreeSurfer subdivisions. Previous work has compared FreeSurfer’s hippocampal subfields to hand drawn volumes (Herten et al., 2019; Worker et al., 2018); however, to our knowledge there is yet to be any comparison of automated amygdala nuclei segmentation to hand-tracing. Reliable methods exist for expert manual segmentation of the amygdala (Aghamohammadi-Sereshki et al., 2018; Entis et al., 2012); however, this typically requires high-resolution and high-field strength neuroimaging (i.e., >3T MRI Scanner, sub-millimeter voxels). Studies looking at the degree of overlap between such methods and the FreeSurfer algorithm for amygdala segmentation would be helpful for effective evaluation of validity.

In considering our results, it is important to note a few potential limitations of our work. First, we processed repeated MRI images using the cross-sectional FreeSurfer pipeline (and not the longitudinal stream of this software suite). This may be a debated choice. However, FreeSurfer’s longitudinal pipeline is not actually independently “considering” (segmenting) the different MRI scans. This may violate some theoretical aspects of test-retest reliability and produce a more favorable set of reliability metrics for FreeSurfer’s methods. Second, we only used T1-weighted scans in FreeSurfer, but additional MRI volumes (e.g., T2-weighted) from the same subjects would likely yield a more reliable segmentation. FreeSurfer’s developers have worked to allow the amygdala and hippocampal subdivision routines to accept high-resolution T2-weighted volumes, and this should be investigated in future work. Third, the sample is a rather homogenous group of individuals and may not represent the greater population. All participants were recruited from the University of Amsterdam, with limited race and ethnic variability. Similarly, all participants were neurotypical young adults in a constrained age range (Mean=22.08+/-1.88). Additional work considering reliability of the method in a diverse set of populations (e.g., pediatric, elderly, mild cognitive impairment) would be helpful in ascertaining how well these findings generalize outside of our sample population.

Limitations notwithstanding, our work extends the information provided by previous publications regarding the reliability of FreeSurfer’s subcortical segmentation for the hippocampus, amygdala, and their respective subregions. To our knowledge, this is the first work to directly investigate the test-retest reliability of the amygdala nuclei algorithm in FreeSurfer 7. The strengths of our work include a reasonably large sample size, the use of FreeSurfer’s more robust longitudinal pipeline, and the report of mathematically rigorous measures of reliability. Our work provides additional confidence in interpreting those regions with high reliability and a necessary caution in interpretation of those with poorer results.

## Data Availability Statement

Neuroimaging data used in our analyses were sourced from the Amsterdam Open MRI Collection (AOMIC, Snoek et al., 2021). FreeSurfer software is publicly and freely available from the FreeSurferWiki resource (http://surfer.nmr.mgh.harvard.edu/fswiki/FreeSurferWiki), which is developed and maintained at the Martinos Center for Biomedical Imaging (http://www.nmr.mgh.harvard.edu/martinos/noFlashHome.php). This software, information and support are provided online at the FreeSurferWiki webpage.

